# Longitudinal Assessment of Fluorescence Stability Shows Fluorescence Intensity Decreases Over Time: Implications for Fluorescence Microscopy Studies

**DOI:** 10.1101/2025.10.15.682696

**Authors:** Sean C. Sweat, Sarah P. R. Berg, Tenzin Kunkhyen, Emma G. Foster, Claire E. J. Cheetham

**Affiliations:** Department of Neurobiology, University of Pittsburgh, Pittsburgh PA 15213; Center for Neuroscience, University of Pittsburgh, Pittsburgh, PA 15213; University of Pittsburgh-Carnegie Mellon University Medical Scientist Training Program, Pittsburgh, PA 15261

## Abstract

Immunohistochemistry (IHC) is one of the most widely used techniques across basic, translational, and clinical sciences. Key considerations need to be made to achieve reliable and robust IHC staining, however what has been understudied is the stability of IHC signal intensity over time. Changes in signal intensity over time have significant implications for data analysis and interpretation and ultimately impact scientific conclusions. In order to explore changes in IHC signal, the stability of fluorescence intensity was assessed over the course of six weeks using widefield or confocal microscopy. Results indicate that fluorescence intensity can decrease over this time course and that whether this decrease occurs and to what extent is influenced by the selection of the primary antibody as well as that of the secondary antibody, primary-secondary antibody combination, and utilization of chemical staining versus IHC staining. This investigation reinforces best practices for imaging fluorescent staining to ensure accurate and reliable data collection, be it for cell counting, assessing protein expression levels, or marker colocalization.

## Introduction

Immunohistochemistry (IHC) is one of the most broadly used techniques in basic, translational, and clinical laboratories. Indeed, a PubMed search indicates that more than 790,000 articles have been published since the 1950s that have either utilized IHC or discussed its principle. It takes advantage of the immunological concept of antibody targeting of specific antigens to label, and thus identify, proteins in fixed tissue that can then be imaged and analyzed. The history of IHC extends as far back as 1941, the year in which Albert Coon and colleagues developed the first direct staining method: using frozen tissue sections, they successfully labeled Type III pneumococci with fluorescent dye-conjugated primary antibodies [1] (for a more extensive history of IHC see [2]). Scientists have since expanded the IHC technique by producing enzyme-labeled primary antibodies [3, 4] and developing chromogenic multiplexing to better capture complex microenvironments in tissue specimens [5]. Fluorescence multiplexing has also become a commonly used approach across multiple disciplines to investigate cellular components in detail. With the widespread use of fluorescent microscopy came also the development of fluorescent dyes that are both more resistant to photobleaching (a phenomenon in which fluorescent signal is gradually lost with repeated illumination) and less phototoxic (damaging to tissue specimens). The formation of reactive oxygen species (ROS) is known to be a cause of photobleaching and phototoxicity. Thus cyanine and rhodamine-based fluorescent dyes have undergone chemical-based improvements such as 1) incorporation of quenchers that eliminate the formation of ROS when fluorophores are illuminated, and 2) addition of chemical groups that either prevent ROS from reaching the fluorophore or prevent the formation of ROS altogether [6].

IHC can be performed on samples that have been processed in various ways, including paraffin-embedded sections (often used in clinical settings), cryo-embedded, or gelatin-embedded sections. Despite the various approaches to IHC, the steps involved are similar across variations of the technique: rehydration of tissue (for paraffin-embedded sections), possible antigen retrieval, tissue blocking, primary antibody incubation, secondary antibody incubation, enzyme-substrate visualization if needed, and counterstaining/coverslipping. Given the series of steps involved, the key considerations to obtain strong and specific staining using IHC include: 1) antibody selection and verification of specificity; 2) optimization of antigen retrieval method (if needed); 3) choice of detection method, be it a fluorescent dye, chromogen, or enzyme; and 4) inclusion of positive and negative controls. Nevertheless, many difficulties can arise with IHC, from absent, weak or patchy staining to significant background fluorescence, yielding a low signal:noise ratio. One important factor that has received little consideration is whether the IHC signal is stable over time after completion of the staining process. In particular, when using fluorescent dyes for antigen detection and quantification, is the fluorescence intensity the same shortly after staining versus several weeks later? If fluorescence intensity changes over time, this could affect detection of antigen labeled cells and structures, quantification of expression levels or colocalization and particularly comparisons between experimental conditions. That is to say, imaging samples shortly after staining versus at a later time point may ultimately influence data interpretation and overall scientific conclusions.

To investigate whether, and to what extent, fluorescence intensity changes over time after completion of IHC staining, olfactory epithelium (OE), olfactory bulb (OB), and whole brain sections were collected from mice, processed with IHC, and imaged weekly over the course of four or six weeks. Various combinations of primary and secondary antibodies were utilized for this investigation to determine whether certain combinations were more susceptible to changes in fluorescence intensity over time. Additionally, the stability of fluorescence intensity from a chemical stain and a fluorescent protein was tested over a six-week time course.

## Methods

### Mouse lines

C57BL/6J (Strain #: 000664), TH-Cre (Strain #: 008601), DAT-Cre (Strain #: 006660), Ai162 (Strain #: 031562), and Ai9 (RCL-tdT) (Strain #: 007909) mice were purchased from The Jackson Laboratory and/or bred in-house. A total of 37 male and female mice were used and maintained in individually ventilated cages at 22 °C and 48% humidity on a 12 h light/dark cycle with ad libitum access to food and water. Mice were group-housed unless same sex littermates were unavailable. Mouse lines used for investigation include DAT-cre;Ai162;Ai9 (for fluorescent protein investigation) and TH-Cre;Ai9 (for anti-TH primary antibody staining).

### Transcardial perfusion, tissue processing, and cryosectioning

Mice were anesthetized with 4% isoflurane in 1L/min O_2_ and transcardially perfused with phosphate buffered saline (PBS) followed by 4% paraformaldehyde (PFA). Heads were removed and post-fixed in 4% PFA overnight, followed by dissection of the target region (olfactory epithelium (OE), olfactory bulb (OB), or whole brain containing cortex and hippocampus) for cryopreservation in 30% sucrose overnight (or until tissue sank). Sections were then embedded in 10% gelatin, fixed/cryopreserved in 2% PFA/15% sucrose overnight, and flash frozen in 2-methylbutane for cryosectioning. OE was coronally sectioned at 50 microns, OB was coronally sectioned at 40 microns, and whole brains for staining of cortex and hippocampus were sagittally sectioned at 40 microns. Sections were stored in PBS or PBS/0.01% sodium azide at 4 °C.

### Immunohistochemistry

The range of time elapsed between mouse perfusion (i.e., tissue fixation) and staining was between one week to five months. For widefield imaging, the time between completion of staining and the first imaging session was a maximum of one week. For confocal microscopy, imaging began within one week of completion of staining, but images from the first two weekly time points could not be used due to technical difficulties with a multi-user instrument. ‘Week 1’ labeled confocal data and images therefore represent data collected less than three weeks after completion of staining. Sections were treated with 1% w/v sodium borohydride in distilled water for 20 minutes, followed by a series of washes in PBS (at least 6 x 3 min) until no residual bubbles were present in solution. Permeabilization and blocking was completed with 5% normal donkey serum (NDS)/0.5% Triton-X100 in PBS for 1 hour. Primary antibody buffer was made with 3% NDS/0.2% Triton-X100 in PBS/0.01% sodium azide. Primary antibodies were added to the buffer at required dilution (see Table 1). The primary antibodies used for this set of experiments were olfactory marker protein (OMP), growth-associated protein-43 (GAP43), hemagglutinin protein (HA), ionized calcium binding adaptor molecule 1 (Iba1), and tyrosine hydroxylase (TH). Primary antibody incubation duration varied depending on antibody (see Table 1), either 2- or 4-day incubations at 4°C, followed by a series of washes (3 x 5 min) in PBS. Secondary antibody buffer was made with 3% NDS/0.2% Triton-X100 in PBS. Secondary antibody was added to buffer and sections were incubated in secondary antibody for 1 hour at room temperature followed by a series of washes (3 x 5 min) in PBS. Sections were mounted onto glass slides, covered with Vectashield containing DAPI, and coverslipped. Nonhardset Vectashield DAPI medium was used for OE sections, whereas Hardset Vectashield DAPI medium was used for OB and cortical/subcortical sections.

**Table 1.** Summary table of primary and secondary antibodies used. Species that antibody was raised in, dilution used, incubation time, manufacturer, and catalog number are provided.

| Antibody [species] (Dilution) | Incubation Time | Manufacturer and Catalog No. |
| --- | --- | --- |
| OMP [goat] (1:5000) | 4 days | Wako #544-10001 |
| GAP43 [rabbit] (1:1000) | 4 days | Novus Biologicals #NB300-143 |
| HA [rabbit] (1:500) | 4 days | Cell Signaling Technology #3724 |
| Iba1 [rabbit] (1:2000) | 2 days | Wako #019-19741 |
| TH [chicken] (1:2500) | 2 days | AvesLabs #TYH-0020 |
| TH [rabbit] (1:2500) | 2 days | Novus Biologicals #NB300-109 |
| AF488 [donkey anti-rabbit] (1:500) | 1 hour | ThermoFisher Scientific #A21206 |
| AF488 [donkey anti-goat] (1:500) | 1 hour | ThermoFisher Scientific #A11055 |
| AF488 [donkey anti-chicken] (1:500) | 1 hour | ThermoFisher Scientific #A78948 |
| AF546 [donkey anti-rabbit] (1:500) | 1 hour | ThermoFisher Scientific #A10040 |
| AF546 [donkey anti-goat] (1:500) | 1 hour | ThermoFisher Scientific #A11056 |
| AF647 [donkey anti-rabbit] (1:500) | 1 hour | ThermoFisher Scientific #A31573 |
| AF647 [donkey anti-goat] (1:500) | 1 hour | Abcam #Ab150135 |

### EdU click chemistry staining

C57BL/6J mice received an intraperitoneal injection of 50 mg/kg 5-ethynyl-2’-deoxyuridine (EdU (#A10044, ThermoFisher)) in sterile saline and were perfused seven days later. For EdU staining, which occurred after OMP and GAP43 IHC staining, sections were permeabilized for 30 min in PBS/0.5% T-X100. Sections were then incubated in 200 mM copper sulfate, 4 mM sulfo-Cyanine5-azide (#A3330, Lumiprobe), and 200 mg/mL ascorbic acid in PBS for 30 min in the dark. Finally, sections were washed 3 x 5 min in PBS/0.5% T-X100 in the dark, then mounted onto glass slides and coverslipped with Nonhardset Vectashield containing DAPI.

### Mounting of sections from mice expressing a fluorescent protein

DAT-cre;Ai162;Ai9 mice were perfused and OB sections were cut as above. These mice express the genetically encoded calcium indicator GCaMP6s selectively in OB dopaminergic interneurons. OB sections were mounted directly onto glass slides and coverslipped with Hardset Vectashield containing DAPI.

### Widefield microscopy

Widefield imaging was performed using a Revolve widefield microscope (Echo). Settings for each data set, including digital haze reduction, gain, exposure time, and LED power, were optimized at the first imaging session, and these identical settings were used for each subsequent imaging session. Of note, specific imaging parameters were applied to each individual section used for analyzing anti-TH primary antibody and GCaMP6s fluorescence intensity, thus a range of parameters is reported in Table 2; however, identical settings were maintained for each weekly imaging session for each individual section. Between imaging sessions, slides were kept covered in a slide box and refrigerated at 4 °C. OE sections were imaged using a 20x objective (Olympus UPlanXApo, NA 0.80), with 2-3 regions per section selected for imaging. These regions were either along the septum of the OE or among the turbinates and could be re-located at each imaging session. OB and whole brain sections were imaged using a 10x objective (Olympus UPlanXApo, NA 0.40). Control experiments were performed using OMP-stained OB sections. Control experiment 1 consisted of imaging once per day for seven days. Control experiment 2 consisted of imaging once, waiting six weeks, then imaging again. Control experiment 3 consisted of imaging every two minutes until seven data points were collected. For all other experiments, images were collected every seven days. For OB and whole brain sections individual tiled images were collected in grayscale tiff format, converted to jpeg files, and then stitched within Affinity Photo software by using the Panorama function. Stitched images were then exported in tiff format. Image brightness was not altered throughout analyses or in manuscript figures.

**Table 2.** Summary of widefield imaging parameters. Combination of primary antibody and fluorophore is provided. Revolve widefield microscope (Echo) imaging parameters are also provided including digital haze reduction, gain, exposure, and LED power.

| Experiment/Analysis | Primary antibody | Fluorophore/ fluorescent protein | Digital haze reduction | Gain | Exposure (ms) | LED power (%) |
| --- | --- | --- | --- | --- | --- | --- |
| OE (single stain) | OMP | AF488 | 3 | High | 30 | 30 |
| OE (single stain) | HA tag | AF546 | 3 | High | 70 | 100 |
| OE (triple stain-widefield) | GAP43 | AF546 | 3 | Low | 90 | 100 |
|  | OMP | AF488 | 3 | Low | 200 | 100 |
|  | EdU | AF647 | 3 | Low | 80 | 100 |
| OB GL (single stain) | OMP | AF488 | 3 | High | 50 | 40 |
| OB GL & GCL (single stain) | GAP43 | AF488 | 3 | High | 35 | 45 |
|  |  | AF546 | 3 | High | 60 | 100 |
|  |  | AF647 | 3 | High | 80 | 100 |
| OB GL (single stain-Chicken Antibody) | TH | AF488 | 3 | Low or High | 50 - 110 | 100 |
| OB GL (single stain- Rabbit Antibody) | TH | AF488 | 3 | Low or High | 80 - 200 | 50 - 100 |
| OB GL GCaMP6s in dopaminergic neurons | N/A | GCaMP6s | 3 | Low | 310 -350 | 60 - 80 |
| Cortex (single stain) | Iba1 | AF546 | 3 | High | 800 | 70 |
| Hippocampus (single stain) | Iba1 | AF546 | 3 | High | 800 | 70 |
| Control Exp. 1 (OB, single stain) | OMP | AF488 | 3 | High | 85 | 80 |
|  |  | AF546 | 3 | High | 170 | 100 |
|  |  | AF647 | 3 | High | 525 | 100 |
| Control Exp. 2 (OB, single stain) | OMP | AF488 | 3 | High | 85 | 80 |
|  |  | AF546 | 3 | High | 170 | 100 |
|  |  | AF647 | 3 | High | 525 | 100 |
| Control Exp. 3 (OB, single stain) | OMP | AF488 | 3 | High | 85 | 80 |
|  |  | AF546 | 3 | High | 170 | 100 |
|  |  | AF647 | 3 | High | 525 | 100 |

### Confocal microscopy

Confocal imaging was performed using an Olympus Fluoview 3000 confocal microscope. A separate set of triple-stained OE sections (GAP43, OMP, EdU) underwent confocal imaging and analyses for five consecutive weeks. Images were collected with a 60x oil immersion objective (Olympus UPlanXApo, NA 1.42), with an image size of 512 x 512 pixels and three adjacent images stitched together. No averaging was used. Settings for each channel, including laser power, gain and sampling speed, were optimized at the first imaging session, and the same settings were used for each subsequent imaging session (Table 3). Step size for z stack collection was 1 μm. Between imaging sessions, slides were kept covered in a slide box and refrigerated at 4 °C.

**Table 3.** Summary of confocal imaging parameters. Combination of primary antibody and secondary antibody fluorophore is provided. Confocal imaging parameters provided, including laser power, gain, sampling speed, and pixel size.

| Primary Stain<br>(Secondary<br>Fluorophore) | Excitation<br>wavelength (nm) | Laser<br>power (%) | Gain | Sampling speed<br>($\mu\text{s}/\text{pixel}$) | Pixel size (xy)<br>($\mu\text{m}/\text{pixel}$) |
| --- | --- | --- | --- | --- | --- |
| OMP (AF488) | 488 | 2.0 | 1x | 2.0 | 0.414 |
| GAP43 (AF546) | 561 | 2.0 | 1x | 2.0 | 0.414 |
| EdU (AF647) | 640 | 4.0 | 1x | 2.0 | 0.414 |

### Image analysis

For OE sections, a region of interest (ROI) was selected from each imaged area for analyses. Two to three OE sections from each mouse were stained, and two to three areas were imaged within each OE section. Within each imaged area one ROI was selected for analyses. The width of these selected regions spanned from the apical surface of OE to the basal lamina. For OB sections, the entire layer that contained stained neurons or axons (e.g. glomerular layer (GL), granule cell layer (GCL), etc.) was used as the ROI for analyses. For OMP- and GAP43-stained OB sections, the channel in which the relevant staining could be observed was used to draw ROIs. For cortex, ROIs within Layer 1 were selected for analyses and for hippocampus, ROIs were selected either in the dentate gyrus or Cornu Ammonis subregions. For measurement of GCaMP6s in OB dopaminergic cells, the DAPI channel was used to draw an ROI around the glomerular layer (GL). For each section, ROIs used for the first imaging session (session 1) were used for analysis of images collected during all subsequent sessions. Raw mean gray fluorescence values were measured for each ROI at each imaged time point.

Total (background + staining) fluorescence and background fluorescence values were measured to accurately report staining fluorescence. Areas of the section used for assessing background fluorescence contained no staining. Staining fluorescence was calculated by subtracting background fluorescence from total (background + staining) fluorescence. Figure 1 shows an example of this workflow, including raw values.

**Figure 1.**
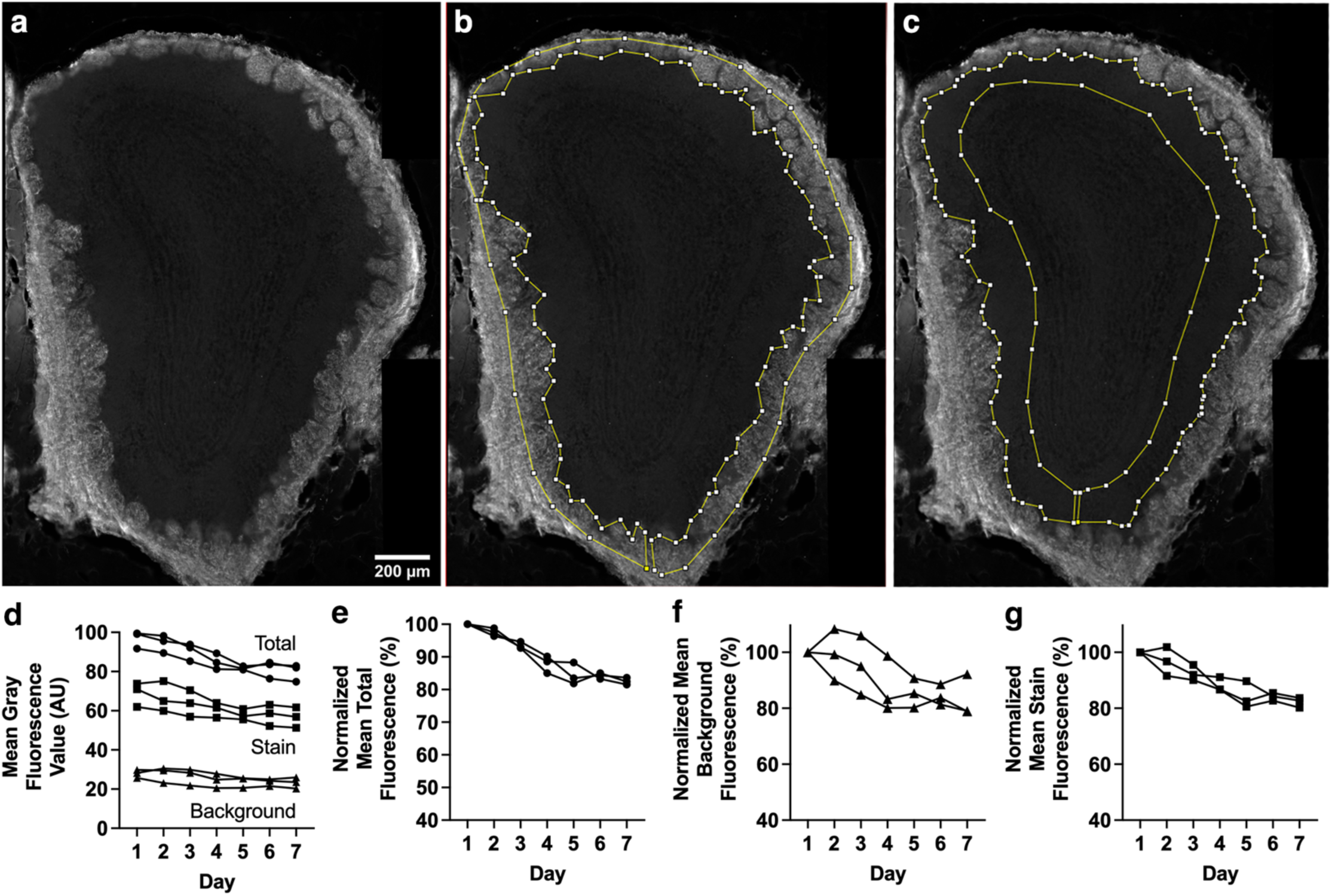
Breakdown of analyses completed for each experiment using Control Exp. 1 (anti-OMP primary, AF488-conjugated secondary) as an example. **a-c.** Example images of OB section used for Control Exp. 1 (anti-OMP-AF488). **a.** Image without overlays. **b.** Same image with overlaid ROI used to assess combined background + staining fluorescence. **c.** Same image with overlaid ROI used to assess background fluorescence (i.e., external plexiform layer (EPL)), in which no OMP staining is present. **d.** Change in raw mean gray fluorescence values for total i.e. background + stain (circle), stain (square), and background (triangle). **e.** Percentage normalized mean gray fluorescence for total (combined background and stain) fluorescence. **f.** Percentage normalized mean gray fluorescence for background fluorescence. **g.** Percentage normalized mean gray fluorescence for staining fluorescence signal (total - background). Each line of data represents a single mouse.

### Statistical Analysis

Staining fluorescence values were normalized as a percentage of the values in session one and further analyzed with GraphPad Prism (version 10.4.2 and 10.5.0) software. The Wilcoxon matched pairs signed rank test was applied to Control Exp. 2. All other experiments underwent one way ANOVA on ranks (Friedman) tests. Two to four mice per experiment were used, with either one to three OB sections per mouse or two to three OE sections per mouse (each OE section had two to three areas imaged and within each imaged area, one ROI was selected for analyses). A summary of statistical results is shown in Table 4.

**Table 4.** Summary of statistical results for each set of analyses comparing fluorescence intensity over time. *= p<0.05; **= p<0.01; ***= p<0.001; NS (non-significant) = p≥0.05.

| Experiment/Analysis (Figure number) | Primary antibody | Fluorophore/ fluorescent protein | Statistical results |
| --- | --- | --- | --- |
| OE (single stain) (Figure 2a-d) | OMP | AF488 | *** |
| OE (single stain) (Figure 2e-h) | HA tag | AF546 | ** |
| OE (triple stain-widefield) (Figure 3a-l) | GAP43 | AF546 | *** |
|  | OMP | AF488 | * |
|  | EdU | AF647 | NS |
| OE (triple stain-confocal, single z plane) (Figure 4d,i,n) | GAP43 | AF546 | *** |
|  | OMP | AF488 | *** |
|  | EdU | AF647 | ** |
| OE (triple stain-confocal, maximum intensity projection) (Figure 4a-c,e,f-h,j,k-m,o) | GAP43 | AF546 | *** |
|  | OMP | AF488 | *** |
|  | EdU | AF647 | ** |
| OB GL (single stain) (Figure 5a-d) | OMP | AF488 | ** |
| OB GL (single stain) (Figure 6a-d, f-i, k-n) | GAP43 | AF488 | ** |
|  |  | AF546 | NS |
|  |  | AF647 | ** |
| OB GCL (single stain) (Figure 6a-c,e,f-h,j,k-m,o) | GAP43 | AF488 | ** |
|  |  | AF546 | NS |
|  |  | AF647 | ** |
| OB GL (single stain-chicken antibody) (Figure 7a-d) | TH | AF488 | ** |
| OB GL (single stain- rabbit antibody) (Figure 7e-h) | TH | AF488 | NS |
| OB GL GCaMP6s in dopaminergic neurons (Figure 7i-l) | N/A | GCaMP6s | * |
| Cortex (single stain) (Figure 8a-d) | Iba1 | AF546 | NS |
| Hippocampus (single stain) (Figure 8e-h) | Iba1 | AF546 | * |
| Control Exp. 1 (OB, single stain) (Figure 9a-l) | OMP | AF488 | ** |
|  |  | AF546 | NS |
|  |  | AF647 | ** |
| Control Exp. 2 (OB, single stain) (Figure 10a-i) | OMP | AF488 | NS |
|  |  | AF546 | NS |
|  |  | AF647 | NS |
| Control Exp. 3 (OB, single stain) (Figure 10j-u) | OMP | AF488 | NS |
|  |  | AF546 | NS |
|  |  | AF647 | NS |

## Results

### Olfactory epithelium sections stained with various antibody combinations show significant decreases in fluorescence intensity over time

Weekly imaging of OE sections stained with anti-OMP primary antibody and AF488-conjugated secondary antibody over a six-week time course showed a significant decrease in mean gray fluorescence (Figure 2a-d; p<0.001), as did sections stained with anti-HA primary antibody and AF546-conjugated secondary antibody (Figure 2e-h; p=0.001). Despite these differing combinations of primary and secondary antibodies both show a decrease in fluorescence intensity, with the anti-HA primary antibody and AF546 combination resulting in fluorescence intensity as low as approximately 30% of the initial value by week 7. This could speak to the importance of stability of the primary antibody used.

**Figure 2.**
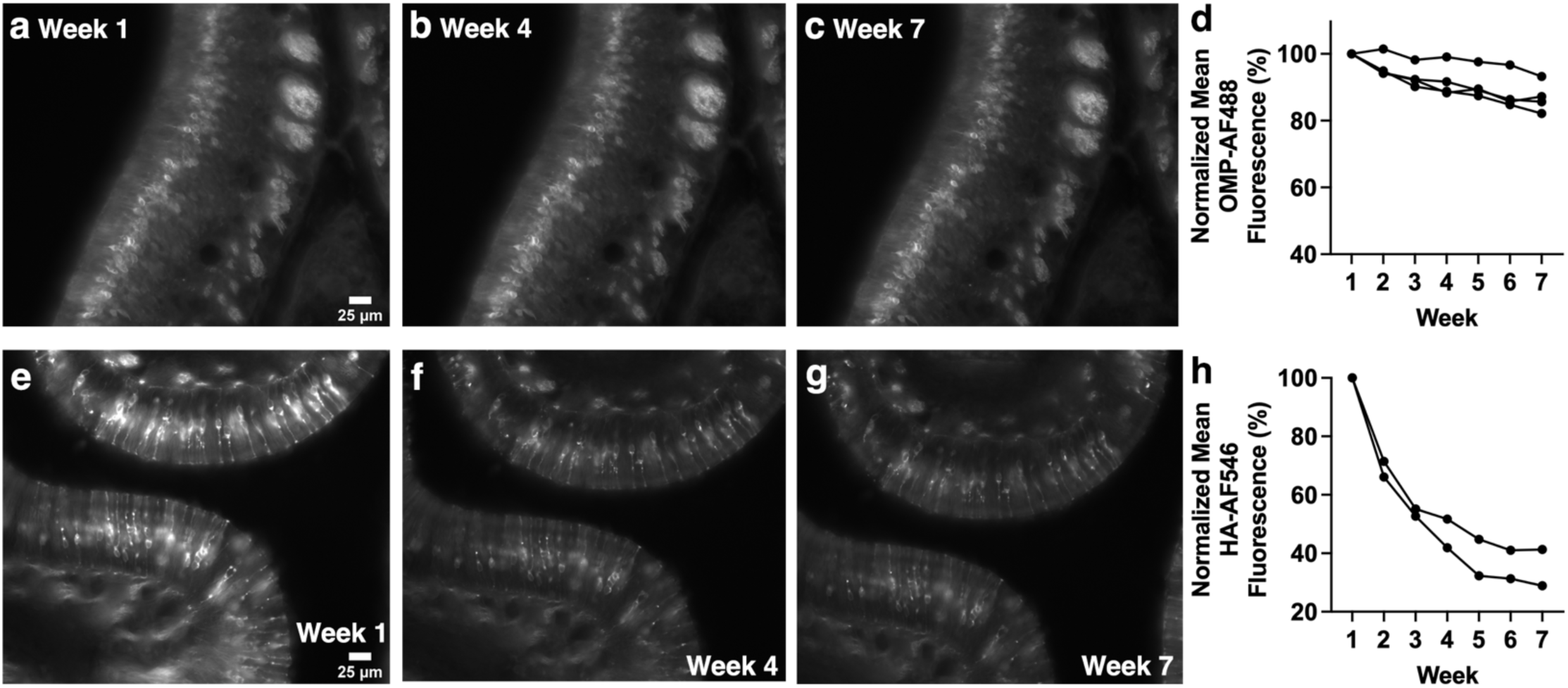
Anti-OMP-AF488 and anti-HA-AF546 stained OE sections show significantly decreased fluorescence intensity over the course of weekly widefield imaging. **a-c.** Anti-OMP primary antibody with AF488 secondary antibody-stained sections imaged with widefield microscopy. **d.** Anti-OMP primary antibody staining combined with AF488-conjugated secondary antibody staining of OE sections show significant decreases in fluorescence intensity. **e-g.** Anti-HA primary antibody-stained with AF546 secondary antibody-stained sections imaged with widefield microscopy. **h.** Anti-HA primary antibody staining combined with AF546-conjugated secondary antibody staining of OE sections show significant decreases in fluorescence intensity. Images of weeks 1, 4 and 7 are shown as examples Each line of data represents a single mouse.

Changes in fluorescence intensity over time were assessed in triple-stained OE sections using both widefield and confocal microscopy. Staining for both the mature OSN marker OMP (AF488 secondary antibody) and the immature olfactory sensory neuron (OSN) marker GAP43 (AF546 secondary antibody) showed significant decreases in fluorescence intensity using widefield imaging (Figure 3a-d, p= 0.033; Figure 3e-h, p<0.001). In contrast, click chemistry staining for EdU (AF647 fluorophore), which birthdates cells born at the time of injection, showed a trend towards a decrease in fluorescence intensity using widefield imaging (Figure 3i-l; p=0.083) that did not reach significance. For confocal images, both single optical sections and maximum intensity projections (MIPs) of z-stacks were analyzed. Both analysis approaches showed significant decreases in fluorescence intensity for OMP/AF488 staining (Figure 4a-e, both p< 0.001), GAP43/AF546 staining (Fig. 4f-j, both p<0.001) and EdU/AF647 (Figure 4k-o, both p=0.003). These results in sum show that triple-stained sections show decreased fluorescence intensity over time, whether imaging with widefield or confocal microscopy.

**Figure 3.**
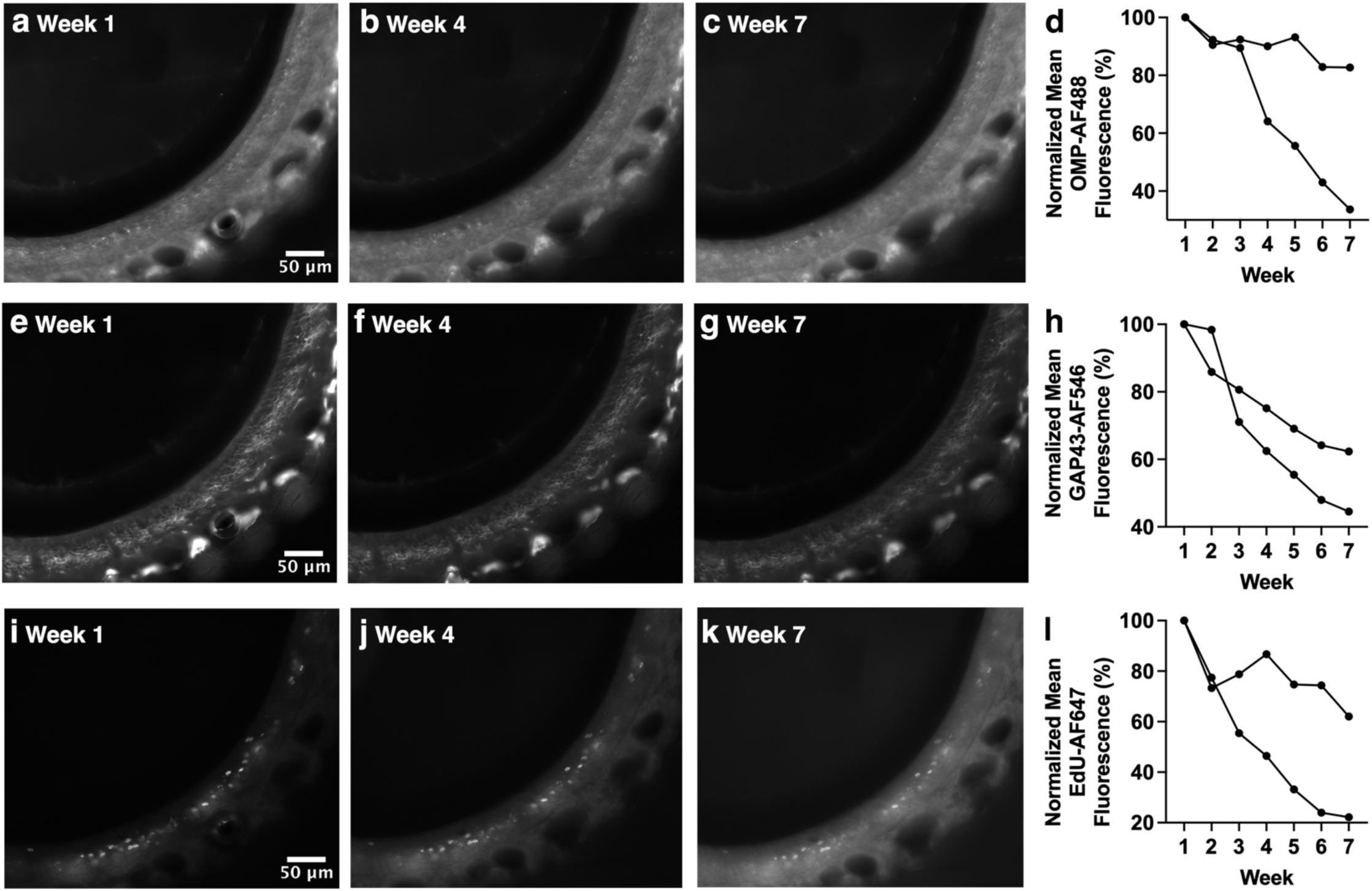
Decreased fluorescence intensity of OE sections triple-stained for OMP-AF488, GAP43-AF546, and EdU-AF647 over the course of weekly widefield fluorescence microscopy. Sections were triple-stained with anti-OMP primary/AF488-conjugated secondary, anti-GAP43 primary/AF546-conjugated secondary and EdU (click chemistry detection with AF647). **a-c** OMP-stained OE section. **d.** Significant decrease in fluorescence intensity for OMP-stained OE over time. **e-g.** GAP43-staining in the same OE section shown in a-c. **h.** Significant decrease in fluorescence intensity for GAP43-stained OE over time. **i-k.** EdU staining in the same OE section shown in a-c and e-g. **l.** Trend towards a decrease in fluorescence for EdU-stained OE over time. Images of week 1, week 4, and week 7 are shown as examples. Each line of data represents an individual mouse.

**Figure 4.**
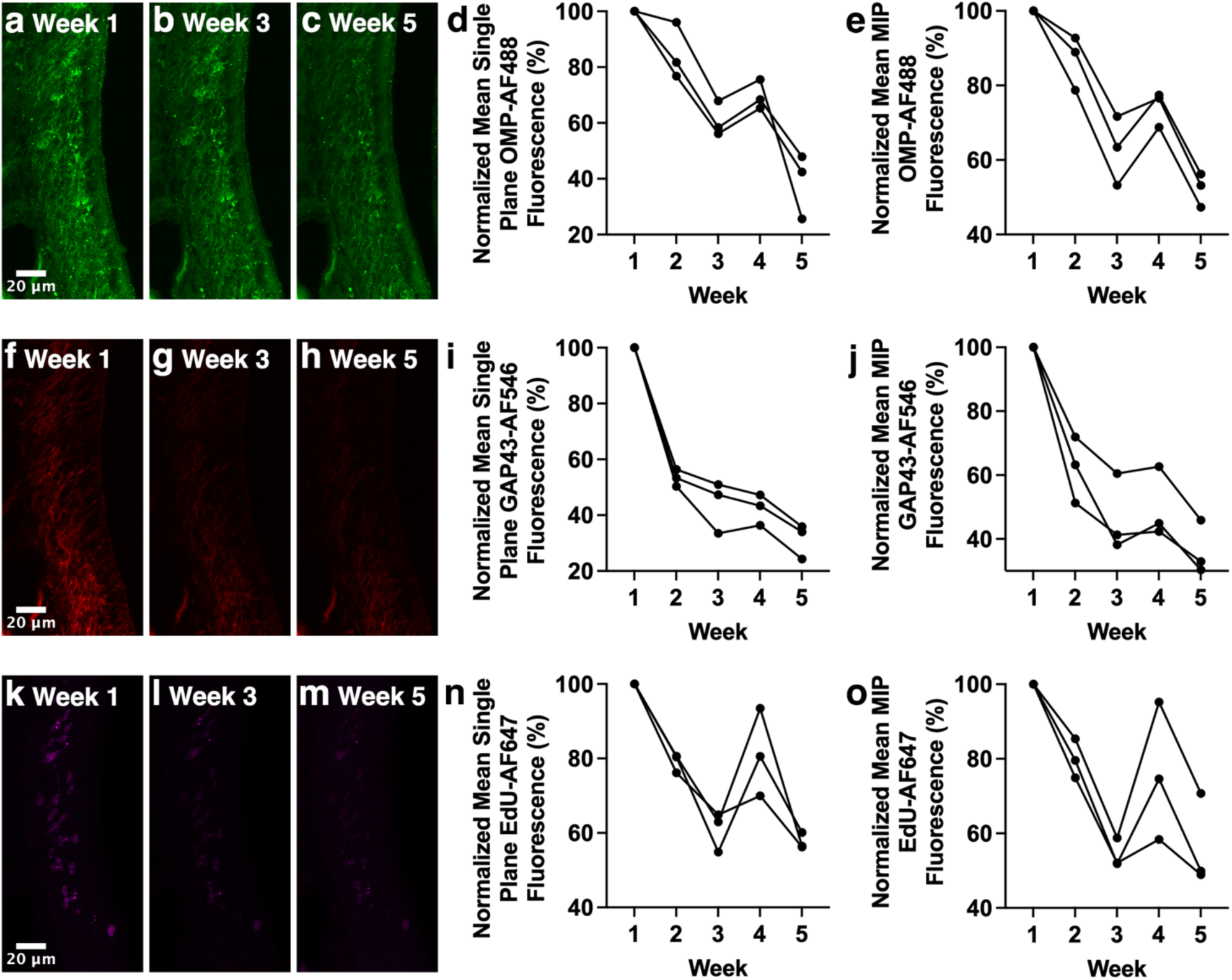
Decreased fluorescence intensity of OE sections triple-stained for OMP-AF488, GAP43-AF546, and EdU-AF647 over the course of weekly confocal microscopy. Sections were triple-stained with anti-OMP primary/AF488-conjugated secondary, anti-GAP43 primary/AF546-conjugated secondary and EdU (click chemistry detection with AF647). **a-c.** MIPs of anti-OMP primary antibody with AF488-conjugated secondary antibody-stained OE sections imaged with confocal microscopy. **d.** Significant decrease in AF488 fluorescence intensity in single optical sections. **e.** Significant decrease in AF488 fluorescence intensity in MIPs. **f-h.** MIPs of anti-GAP43 primary antibody with AF546-conjugated secondary antibody-stained OE sections imaged with confocal microscopy. **i.** Significant decrease in AF546 fluorescence intensity in single optical sections. **j.** Significant decrease in AF546 fluorescence intensity in MIPs. **k-m.** MIPs of EdU with AF647 fluorophore-stained OE sections imaged with confocal microscopy. **n.** Significant decrease in AF647 fluorescence intensity in single optical sections. **o.** Significant decrease in AF647 fluorescence intensity in MIPs. Images of weeks 1, 3 and 5 are shown as examples. Each line of data represents an individual mouse.

### The stability of fluorescence staining intensity in olfactory bulb sections depends on secondary antibody fluorophore

OB sections were imaged weekly. Staining with anti-OMP primary antibody and AF488-conjugated secondary antibody showed a significant decrease in fluorescence intensity across the six weeks of imaging (Figure 5; p=0.007).

**Figure 5.**
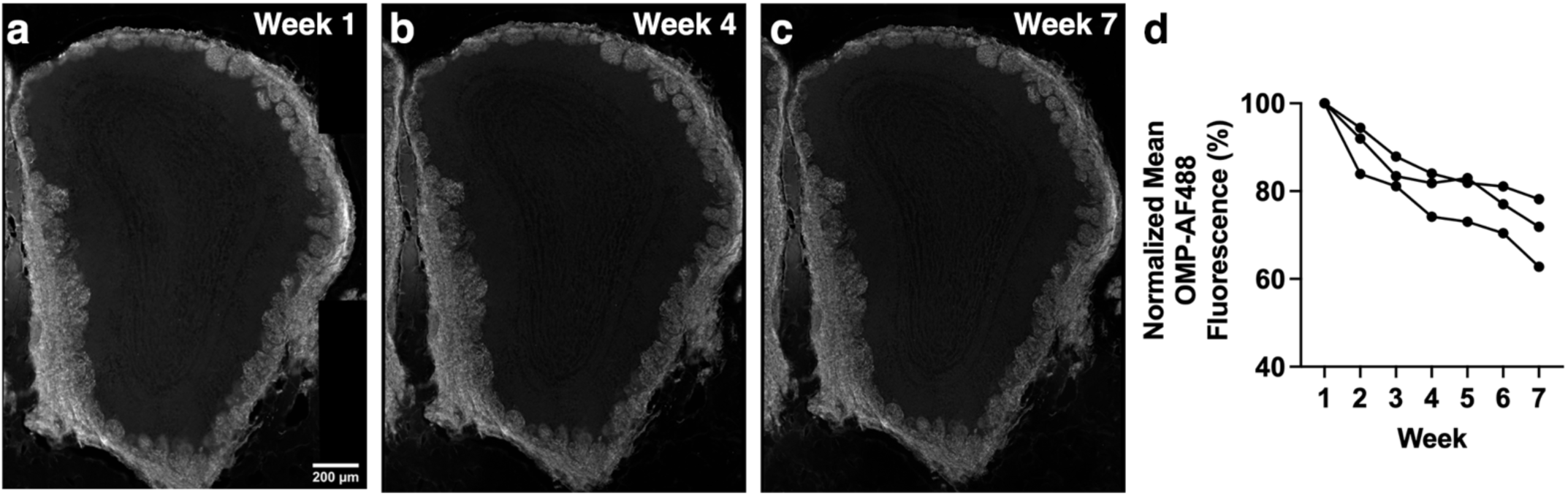
OMP-AF488 stained OB sections showed a significant decrease in fluorescence intensity over time. **a-c.** Anti-OMP primary antibody and AF488-conjugated secondary antibody-stained OB section imaged with wide_b_fi_)_eld microscopy. **d.** OB section imaged weekly show a significant decrease in fluorescence intensity. Images of weeks 1, 4 and 7 are shown as examples. Each line of data represents a single mouse.

Subregions of the OB (both the GL and the GCL) stained for GAP43 were also assessed weekly. For the GL, staining with anti-GAP43 primary antibody and AF488-conjugated secondary antibody staining (Figure 6a-d; p=0.007) or AF647-conjugated secondary antibody staining (Figure 6k-n; p=0.007) showed a significant decrease in fluorescence intensity over the six-week time course. In contrast, anti-GAP43 primary antibody with AF546-conjugated secondary antibody staining showed no significant decrease in fluorescence intensity (Figure 6f-i; p=0.11). Similar results were found in the GCL, with the combination of anti-GAP43 and AF488 staining (Figure 6e; p=0.010) and the combination of anti-GAP43 and AF647 staining (Figure 6o; p=0.007) showing a significant decrease in fluorescence intensity over time, while a combination of anti-GAP43 and AF546 staining did not show a significant decrease (Figure 6j; p=0.11). Regardless of OB subregion, analyzed results were similar. Of additional note is the more gradual decline in fluorescence intensity for anti-GAP43-AF488 staining compared to the large decline in fluorescence intensity observed as early as week 2 for anti-GAP43-AF647 staining.

**Figure 6.**
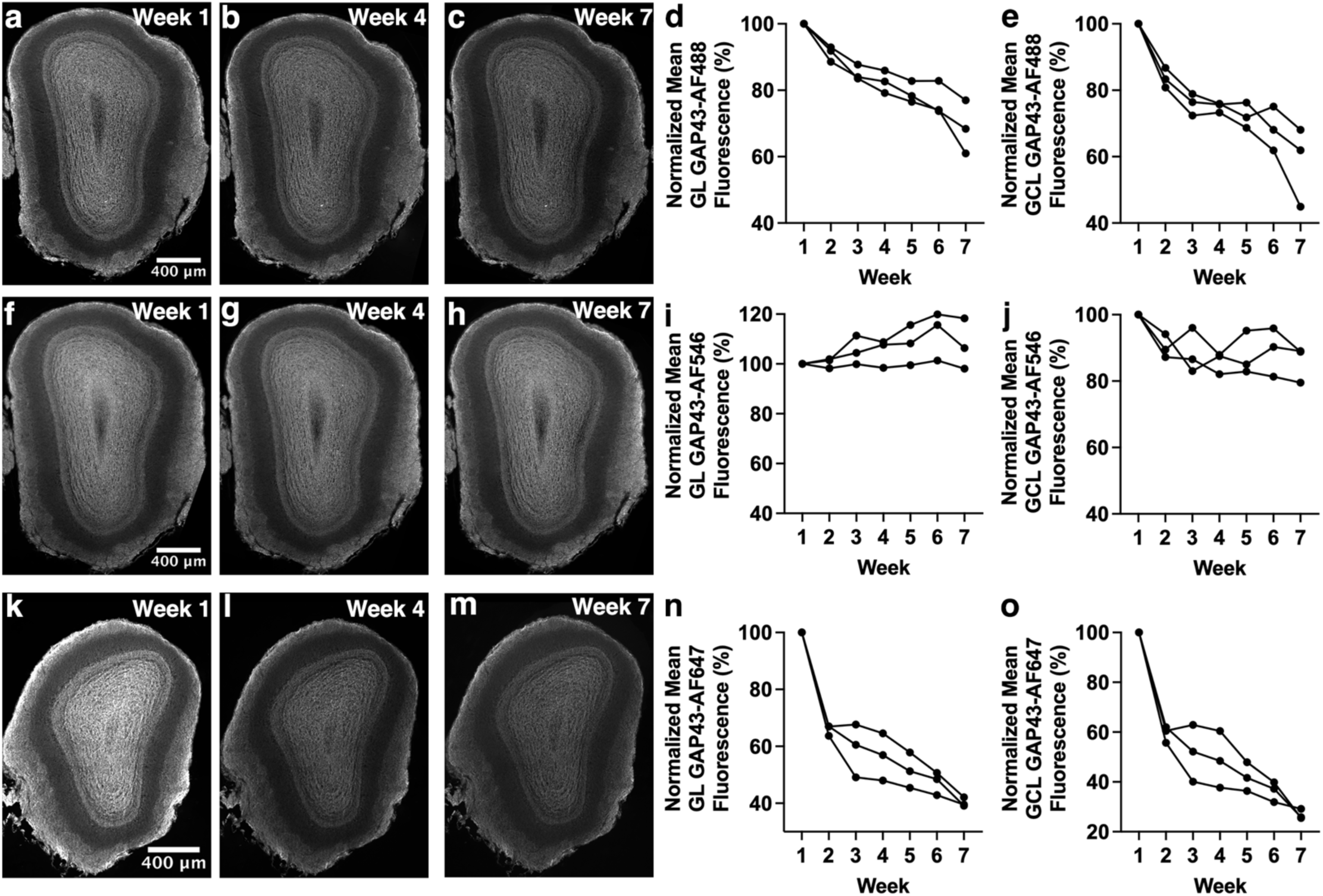
Decrease in fluorescence intensity over time is dependent on secondary antibody fluorophore. **a-c.** Anti-GAP43 primary antibody and AF488-conjugated secondary antibody-stained OB section imaged with widefield microscopy. **d.** Anti-GAP43-AF488 staining in GL shows a significant decrease in fluorescence intensity. **e.** Anti-GAP43-AF488 staining in GCL shows a significant decrease in fluorescence intensity. **f-h.** Anti-GAP43 primary antibody and AF546-conjugated secondary antibody-stained OB sections imaged with widefield microscopy. **i.** Anti-GAP43-AF546 staining in GL shows no significant change in fluorescence intensity. **j.** Anti-GAP43-AF546 staining in GCL shows no significant change in fluorescence intensity. **k-m.** Anti-GAP43 primary antibody and AF647-conjugated secondary antibody-stained OB sections imaged with widefield microscopy. **n.** Anti-GAP43-AF647 staining in GCL shows a significant decrease in fluorescence intensity. **o.** Anti-GAP43-AF647 staining in GCL shows a significant decrease in fluorescence intensity. Images from weeks 1, 4 and 7 are shown as examples. Each line represents data from a single mouse.

### Changes in fluorescence intensity in OB dopaminergic neurons

OB sections were stained to detect TH, which is produced by OB dopaminergic neurons, using two different primary antibodies, raised in either chicken or rabbit. Sections were then incubated with anti-chicken or anti-rabbit AF488-conjugated secondary antibodies and imaged. There was a significant decrease in fluorescence intensity for sections stained with the chicken anti-TH antibody (Figure 7a-d; p=0.001) but only a trend towards a decrease for the rabbit anti-TH antibody (Figure 7e-h; p=0.069) that did not reach statistical significance. This finding indicates that the choice of primary antibody used to detect a particular antigen can influence changes in fluorescence intensity over time. We next assessed whether fluorescence intensity also changes over time for a fluorescent protein. The genetically encoded calcium indicator GCaMP6s, which emits green fluorescence in the presence of calcium and exhibits some baseline fluorescence, was expressed in OB dopaminergic neurons using the DAT-cre;Ai162;Ai9 mouse line (see Methods). GCaMP6s fluorescence intensity in OB dopaminergic neurons also showed a significant decrease across the six-week time course (Figure 7i-l; p=0.017). These results suggest that even fluorescent proteins in fixed tissue undergo changes in fluorescence intensity over time, similar to fluorescent dyes conjugated to antibodies.

**Figure 7.**
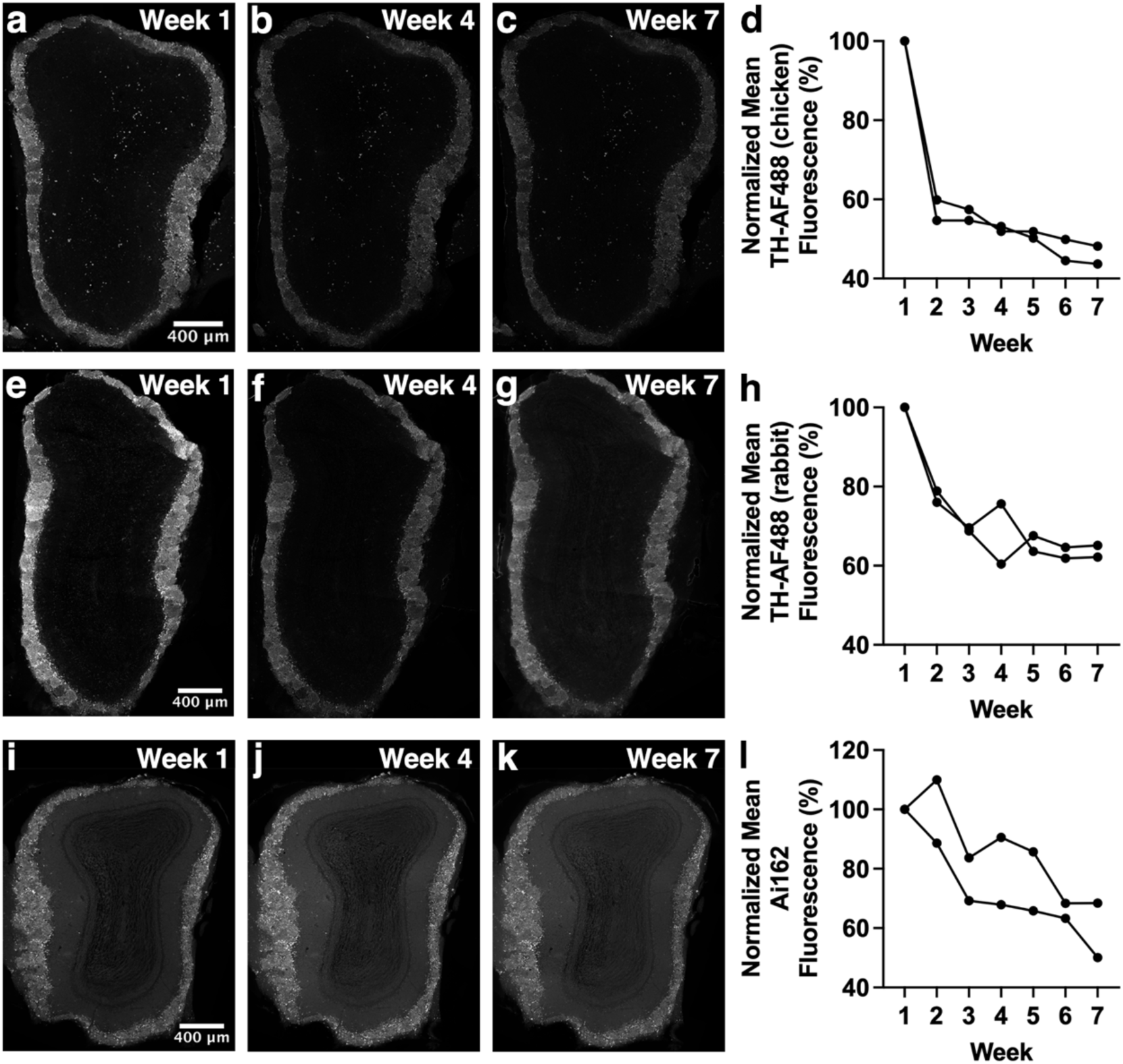
Fluorescence intensity changes over time in OB dopaminergic neurons. **a-c.** Chicken anti-TH primary antibody and AF488-conjugated secondary antibody-stained OB sections imaged with widefield microscopy. **d.** OB sections stained with chicken anti-TH primary antibody and AF488-conjugated secondary antibody show a significant decrease in fluorescence intensity. **e-g.** Rabbit anti-TH primary antibody and AF488-conjugated secondary antibody-stained OB sections imaged with widefield microscopy. **h.** OB sections stained with rabbit anti-TH primary antibody and AF488-conjugated secondary antibody show a trend towards a decrease in fluorescence intensity that does not reach significance. **i-k.** GCaMP6s fluorescence in OB dopaminergic neurons over time imaged with widefield microscopy. **l.** OB sections imaged weekly for GCaMP6s fluorescence show significant decrease in fluorescence intensity. Images of weeks 1, 4 and 7 are shown as examples. Each line of data represents a single mouse.

### Non-olfactory brain regions also show changes in fluorescence intensity over time

Changes in fluorescence intensity were assessed in non-olfactory brain regions, specifically layer 1 of the cortex and dentate gyrus and Cornu Ammonis 1 subregions of the hippocampus. These sections were stained with anti-Iba1 primary antibody, which is a widely used microglial marker, as well as an AF546-conjugated secondary antibody. Cortical sections did not show a significant decrease in fluorescence intensity, however there was a strong trend towards a decrease (Figure 8a-d; p=0.056). As for hippocampus sections, fluorescence intensity decreased significantly (Figure 8e-h; p=0.024). Such results indicate that changes in fluorescence intensity are not restricted to olfactory regions but are also observable in other brain regions, indicating the significance of this phenomenon for labs that utilize IHC regardless of region focus.

**Figure 8.**
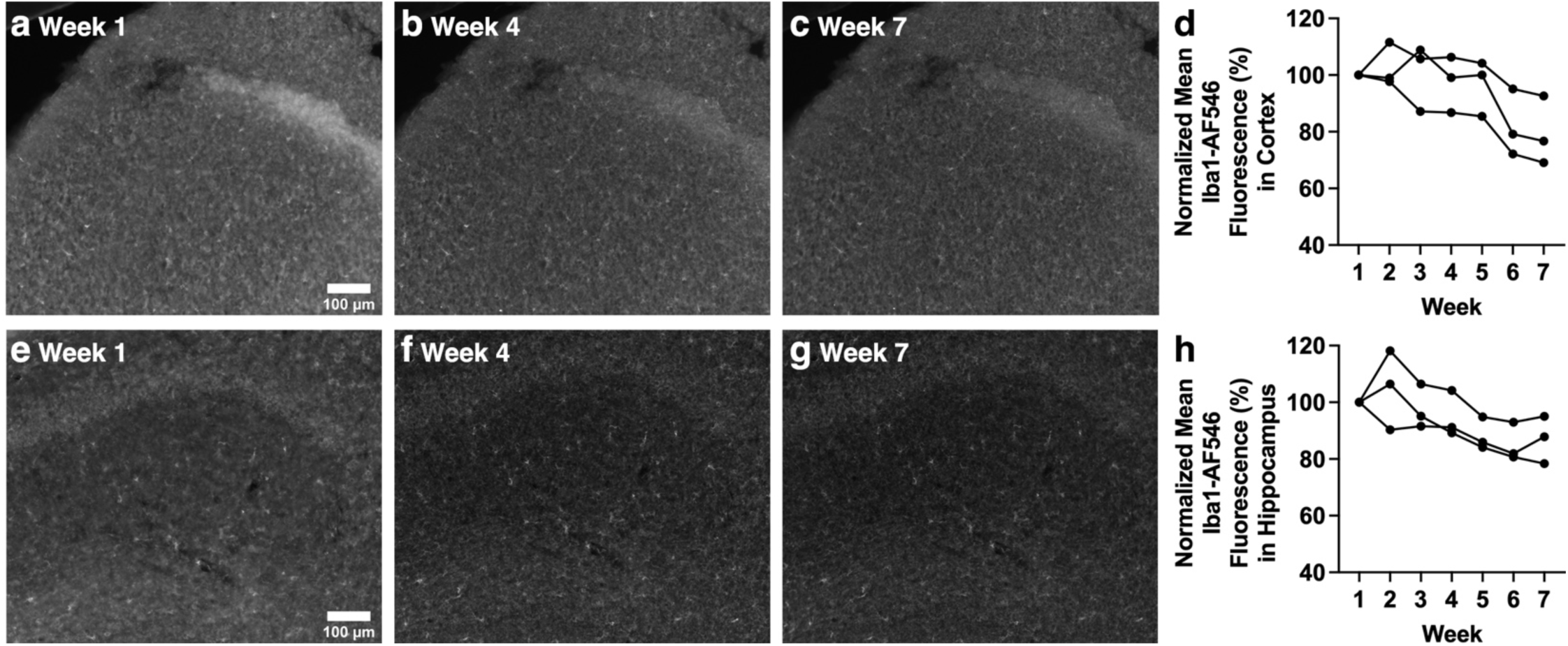
Fluorescence intensity of Iba1 staining in cortex and hippocampus decreases over time. **a-c.** Anti-Iba1 primary antibody and AF546-conjugated secondary antibody-stained cortex layer 1 imaged with widefield microscopy. **d.** Cortical sections stained with Iba1 primary antibody and AF546-conjugated secondary antibody show a trend towards a significant decrease in fluorescence intensity. **e-g.** Anti-Iba1 primary antibody and AF546 conjugated secondary antibody-stained hippocampal dentate gyrus and Cornu Ammonis 1 subregions imaged with widefield microscopy. **h.** Results for hippocampal sections stained with Iba1 primary antibody and AF546-conjugated secondary antibody show a significant decrease in fluorescence intensity. Images of weeks 1, 4 and 7 are shown as examples. Each line of data represents a single mouse.

### AF546-conjugated secondary antibody is most resistant to decreases in fluorescence intensity over time and photobleaching does not underlie the observed changes in fluorescence intensity

To rule out other possible explanations for the patterns of fluorescence intensity changes described earlier, a series of control experiments were carried out. OB sections stained with anti-OMP primary antibody underwent several control experiments, the first being daily widefield microscope imaging for 7 days (Control Exp. 1). Sections stained with an AF488-conjugated secondary antibody showed a significant decrease in fluorescence intensity over the course of daily imaging (Figure 9a-d; p=0.009), as did sections stained with an AF647-conjugated secondary antibody (Figure 9i-l; p=0.006). In contrast, sections stained with an AF546-conjugated secondary antibody did not decrease in fluorescence intensity with daily imaging (Figure 9e-h; p=0.54). A second control experiment was carried out in which sections underwent an initial imaging session, followed by a second and final imaging session six weeks later (Control Exp. 2). AF488 (Figure 10a-c), AF546 (Figure 10d-f), and AF647 (Figure 10g-i) stained sections showed no significant difference in mean gray fluorescence (all p=0.25); however, the small sample size (n=3) combined with only two time points is likely to have precluded detection of significant differences. It is notable that fluorescence intensity for AF488-stained sections decreased by 31% (Figure 10c) and for AF647-stained sections decreased by 78% (Figure 10i), whereas sections stained with AF546-conjugated secondary antibody showed no change in fluorescence intensity at week 7 (Figure 10f). Finally, a third control experiment was carried out in which sections underwent imaging every two minutes (Control Exp. 3), to rule out photobleaching as a possible factor in changes to fluorescence intensity observed in earlier experiments. AF488 (Figure 10j-m), AF546 (Figure 10n-q), and AF647 (Figure 10r-u) stained sections showed no significant difference in fluorescence intensity for this control experiment (p=0.94, p=0.14, p=0.11, respectively), ruling out photobleaching as a possible explanation for results observed in other experiments. It is clear from control experiments 1 and 2 that AF488- and AF647-conjugated secondary antibodies are more prone to decreased fluorescence intensity over time compared to AF546-conjugated secondary antibodies, with AF546-conjugated secondary antibodies maintaining approximately 100% of their original fluorescence intensity. AF647 shows the largest decrease in fluorescence intensity, dropping to as low as 40% of the initial value in Control Exp. 1 and 20% in Control Exp. 2.

**Figure 9.**
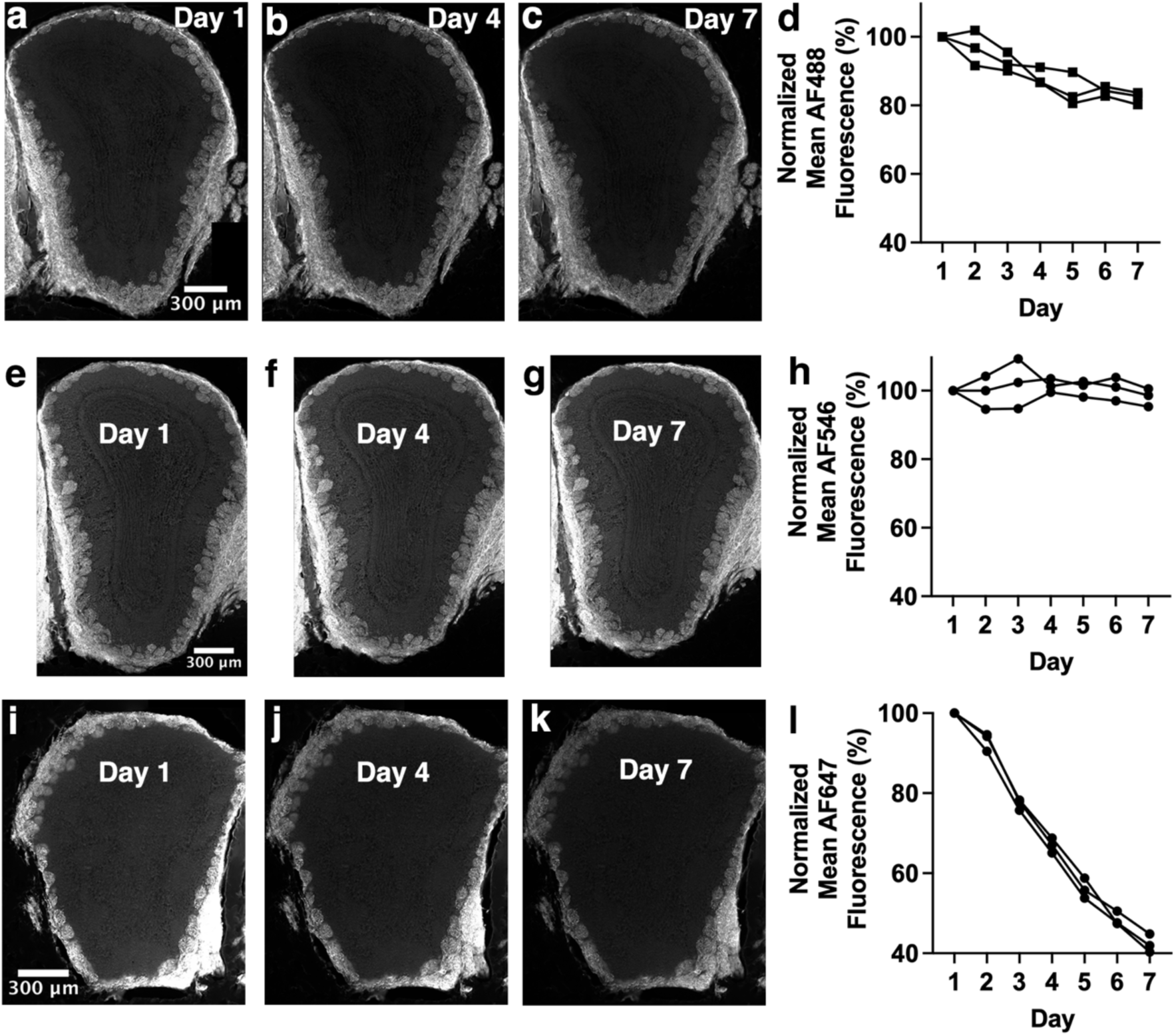
AF546-conjugated secondary antibody is most resistant to decreases in fluorescence intensity over time while AF647-conjugated secondary antibody is least resistant. Control Experiment 1: all sections were stained with anti-OMP primary antibody, followed by staining with AF488-, AF546- or AF647-conjugated secondary antibody and underwent widefield imaging daily for a week. **a-c.** Images of AF488-conjugated secondary antibody-stained sections. **d.** Significant decrease in fluorescence intensity across one week of daily imaging for AF488-conjugated secondary antibody. Note that these are the same data as shown in Figure 1g. **e-g.** Images of AF546-conjugated secondary antibody-stained sections. **h.** No change in fluorescence intensity across one week of daily imaging for AF546-conjugated secondary antibody. **i-k.** Images of AF647-conjugated secondary antibody-stained sections. **l.** Significant decrease in fluorescence intensity across one week of daily imaging for AF647-conjugated secondary antibody. Images of days 1, 4 and 7 are shown as examples. Each line of data represents a single mouse.

**Figure 10.**
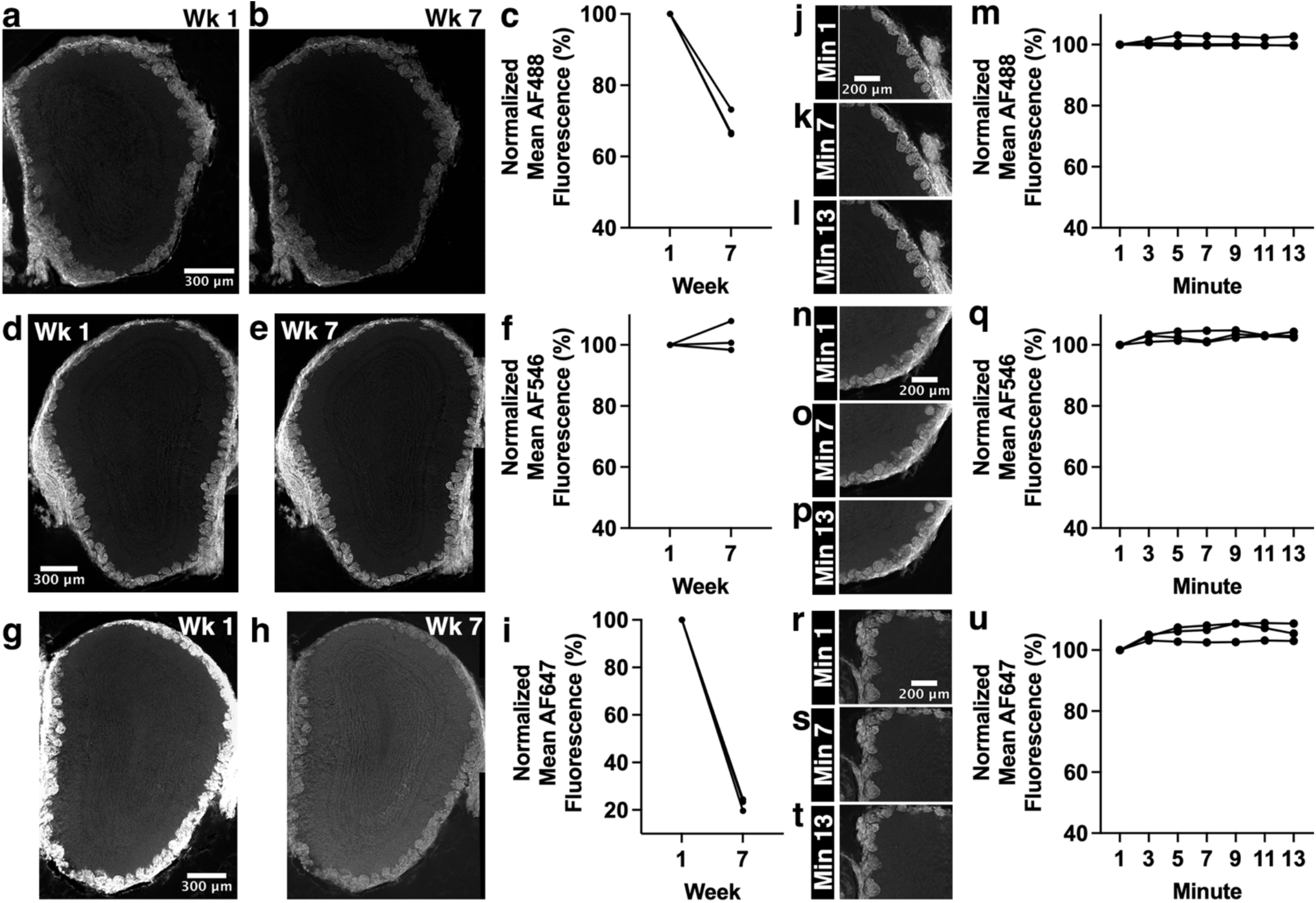
AF546 is the most resistant to fluorescence intensity changes over time and photobleaching does not underlie the observed changes in fluorescence intensity. All sections were stained with anti-OMP primary antibody, followed by staining with AF488-, AF546- or AF647-conjugated secondary antibody. Control Experiment 2 (a-i) consisted of an initial imaging session followed by a second imaging session 7 weeks later, using widefield microscopy. Note that some data points have similar values and overlap (n=3). **a,b.** Images of AF488-conjugated secondary antibody-stained sections at weeks 1 and 7. **c.** No significant difference in fluorescence intensity after six weeks without imaging for AF488-conjugated secondary antibody-stained sections. **d,e.** Images of AF546-conjugated secondary antibody-stained sections at weeks 1 and 7. **f.** No significant difference in fluorescence intensity after six weeks without imaging for AF546-conjugated secondary antibody-stained sections. **g,h.** Images of AF647-conjugated secondary antibody-stained sections at weeks 1 and 7. **i.** No significant difference in fluorescence intensity after six weeks without imaging for AF647-conjugated secondary antibody-stained sections. Control Experiment 3 (j-u) consisted of widefield imaging every two minutes, collecting a total of 7 data points across 13 minutes. **J-l.** Images of AF488-conjugated secondary antibody-stained sections at minutes 1, 7 and 13. **m.** No change in fluorescence intensity for AF488-conjugated secondary antibody-stained sections imaged every two minutes. **n-p.** Images of AF546-conjugated secondary antibody-stained sections at minutes 1, 7 and 13. **q.** No change in fluorescence intensity for AF546-conjugated secondary antibody-stained sections imaged every two minutes. **r-t.** Images of AF647-conjugated secondary antibody-stained sections at minutes 1, 7 and 13. **u.** No change in fluorescence intensity for AF647-conjugated secondary antibody-stained sections imaged every two minutes. Each line of data represents a single mouse.

## Discussion

This study quantified changes in fluorescence intensity over several weeks, which to our knowledge has not been done previously. Using both mouse olfactory and non-olfactory sections, we assessed changes in fluorescence intensity over time with various combinations of primary and fluorescent dye-conjugated secondary antibodies. We found significant decreases in fluorescence intensity over the six-week interval between the first and last imaging session for some antibody combinations and the fluorescent protein assessed. This finding has important implications for accurate interpretation of IHC fluorescence data.

### Secondary antibody dye choice appears to be critical for fluorescence intensity stability

Control experiments using OB sections stained with anti-OMP primary antibody and either AF488-, AF546-, or AF647-conjugated secondary antibodies provided valuable insight into changes in fluorescence intensity over time. Control Exp. 1, in which sections were imaged once a day for 7 days, showed AF546 to be most resistant to changes in fluorescence intensity during this time frame (Figure 9h, 1.86% decrease), with AF488 being somewhat susceptible (Figure 9d, 17.8% decrease), and AF647 being highly susceptible (Figure 9l, 57.7% decrease). In control Exp. 2, in which sections were imaged only twice, at week 1 and week 7 time points, AF546 was again the most resistant to changes in fluorescence intensity even over this 7-week time course (Figure 10f). Control Exp. 3, in which sections were imaged every other minute during a single imaging session, showed all three dyes to have maintained stable fluorescence intensity (Figure 10m,q,u), ruling out photobleaching as a possible explanation for changes in fluorescence intensity over longer time courses in other experiments.

This trend in resistance to decreases in fluorescence intensity, with AF546 being the most resistant, AF488 being somewhat susceptible, and AF647 being the most susceptible, was also observed in imaging the GL and GCL subregions of the OB, in which anti-GAP43 primary antibody was used. AF546 stained sections showed no significant difference in fluorescence intensity over time (Figure 6i,j), whereas AF488-stained sections and AF647-stained sections showed a significant reduction in fluorescence intensity (Figure 6d-e,n-o). AF488 stained sections showed a more gradual decrease in fluorescence intensity whereas AF647-stained sections showed a rapid decrease in fluorescence intensity, reducing by as much as ∼40% from week 1 to week 2. Interestingly, OE sections stained with the anti-HA primary antibody and AF546-conjugated secondary antibody did show a significant decrease in fluorescence intensity over the course of seven imaging sessions (Figure 2e-h). It is possible that this particular combination of primary and secondary antibodies is especially sensitive to changes in fluorescence over time. Additionally, the combination of GAP43 primary antibody and AF546-conjugated secondary antibody staining decreased significantly in fluorescence intensity in triple-stained sections, whether imaged with widefield or confocal microscopy (Figure 3e-h,l; Figure 4f-j). In this instance, the AF546 resistance to fading fluorescence may be lost as a result of staining for multiple antigens in a single section and/or the result of combining this antibody staining with a chemical stain (EdU). Hippocampal sections stained with a combination of anti-Iba1 primary antibody and AF546-conjugated secondary antibody also showed a significant decrease in fluorescence intensity over time (Figure 8h); however, this decrease is smaller in magnitude than that of HA-stained OE or triple-stained OE sections. This could speak to the importance of considering tissue type and/or differences in staining brain regions of various cytoarchitectures.

### Chemical staining shows a different pattern of fluorescence intensity decrease compared to IHC staining

OE sections were triple-stained with the following combinations: GAP43-AF546, OMP-AF488, EdU-AF647. Interestingly, EdU chemical staining combined with AF647 fluorophore in these sections showed a different pattern of fluorescence intensity change compared to other sections stained with AF647 in this investigation (Figure 3i-l; Figure 4k-o). Indeed, the decreases in EdU-AF647 fluorescence intensity was not as rapid compared to GAP43-AF647 staining in GL and GCL OB subregions (Figure 6n-o). Between week 1 and week 2, GAP43-AF647 stained sections showed a decrease in fluorescence intensity of ∼40%, whereas EdU-AF647 fluorescence intensity decreased ∼25% during this time period (with widefield imaging). Hence, the same fluorescent dye may behave differently over time depending on staining modality and/or whether the dye is antibody-conjugated or not.

### Similar patterns of changes in fluorescence intensity are observed whether utilizing widefield or confocal microscopy

Sets of triple-stained OE sections underwent both weekly widefield imaging and weekly confocal imaging (using different sets of sections, with analyses of single optical sections and MIPs for confocal images). GAP43-AF546 staining within these triple-stained sections showed similar changes in fluorescence intensity across imaging and analysis approach (Figure 3d; Figure 4i-j), indicating that for this antibody combination, widefield and confocal imaging report similar changes on fluorescence intensity.

As for OMP-AF488 and EdU-AF647 staining (Figure 3a-d,i-l; Figure 4a-e,k-o) widefield imaging results showed a consistent decrease in fluorescence intensity from week to week and confocal imaging showed a decrease in fluorescence intensity that eventually slowed. These differences between widefield and confocal imaging could be due to fundamental differences in illumination mode. Widefield microscopy results in illumination of areas of the section that are outside the focal plane whereas confocal microscopy involves illuminating a particular focal point at any given time. Additionally, the laser light source in confocal microscopy versus the LED light source in widefield microscopy could contribute to the different patterns of fluorescence intensity changes observed.

Overall, these results indicate that whether one is using widefield imaging or confocal imaging approaches, changes in fluorescence intensity occurs, over a time course of weeks, and should be taken into consideration when designing experimental and analytical approaches. Given that our results support that changes in fluorescence do not appear to be specific to imaging modality, one would expect such changes in fluorescence intensity to be observed with other imaging modalities as well.

### Reduced fluorescence intensity over time also occurs in non-olfactory brain regions

Several non-olfactory sections were stained with Iba1 and AF546-conjuagated secondary antibody in order to investigate changes in fluorescence intensity in other brain regions (Figure 8). Cortical sections show a strong trend towards decreased fluorescence intensity over time (p=0.056), and hippocampal sections showed a significant decrease. These results support that fluorescent proteins and both olfactory and non-olfactory sections undergo changes in fluorescence intensity.

### Possible contributors to decreased fluorescence intensity over time

Given the result of the AF546 fluorophore showing greater resistance to decreases in fluorescence intensity over time compared to AF488 or AF647 in a number of experiments, one important consideration is the differences in chemical structure of each fluorophore [7, 8]. Such differences in chemical structure would imply different chemical interactions with the local microenvironment, i.e., interaction of the fluorophore with local proteins or DNA structures (in the context of nuclear staining). It is possible that the types of chemical interactions that a fluorophore has with the targeted protein/DNA structure of interest or with other proteins/structures in the local environment can influence its photostability over time. This could indeed explain why the combination of anti-HA primary antibody with AF546 and the AF546 fluorophore in triple-stained sections did not show the same resistance to decreased fluorescence intensity compared to the combination of GAP43 and AF546 staining in OB sections. Interaction of AF546 with the anti-HA primary antibody or with the HA protein target may be different from the interaction of AF546 with the anti-GAP43 primary antibody or with the GAP43 protein target, leading to differences in photostability. Triple-stained OE sections were stained with the following primary and secondary staining combinations: anti-GAP43 primary-AF546-conjugated secondary; anti-OMP primary-AF488-conjugated secondary; and EdU-AF647. Thus, AF546 fluorophores in triple stained sections may interact with AF488 and AF647 fluorophores present in the section as well as with anti-OMP primary antibodies and EdU staining, ultimately affecting their photostability. Additionally, fluorophores are susceptible to self-quenching, a phenomenon in which fluorophores, when in close proximity, can interact and thus reduce fluorescence intensity as a result of energy transfer and dimerization [9]. Self-quenching is more likely to occur when there is a high concentration of the fluorophore. Given results collected from this investigation it is possible that AF488, AF546, and AF647 are susceptible to self-quenching at different rates.

As for decreased fluorescence intensity of GCaMP6s, a key consideration is its structural makeup, which includes a circularly permuted Green Fluorescent Protein (GFP), calmodulin, and an M13 fragment of the myosin light chain kinase. It is worth speculating that protein-protein interactions of the GFP chromophore with calmodulin and the M13 fragment can influence its photostability over time.

Lastly, the type of antifade reagent utilized could also be a significant contributor – for this investigation both Hardset and NonHardset VectaShield mounting medium were used, but it is possible that other mounting medium types could have differing effects on fluorescence intensity over time. Indeed, the original study that introduced Alexa fluorophores presented evidence that fluorophore brightness and photostability can be influenced by the chosen mounting medium [7]. Furthermore, another study showed that Vectashield mounting medium can contribute to reduced AF647 fluorescence [10].

### Limitations and future directions

This investigation involved imaging over a four-week (in the case of confocal imaging of triple stained OE sections) or six-week time period. It is possible that further changes in fluorescence intensity would be noted at even later time points, such as several months after the initial imaging session. This would provide additional insight into how long after the initial imaging session fluorescence intensity may continue to change, i.e., would fluorescence intensity signal eventually reach 0% if assessed at a late enough time point?

Also of note, is the primary use of olfactory tissues in this investigation. Although non-olfactory regions, including the cortex and hippocampus, were used, other brain regions should be assessed given that differences in brain cytoarchitecture could lead to different patterns of fluorescence intensity changes over time. An even broader assessment of other tissues throughout the body would prove informative as well, including embryonic tissue or tissue from other organs such as liver or heart.

Moreover, not all possible primary antibody-secondary antibody combinations relevant to the olfactory system were exhausted for this investigation. Additionally, assessing other types of fluorophores, as well as chromophores and chromogens, would be of value. Of note is the use of Alexa dyes for this investigation–different results might be observed if other types of fluorescent dyes are utilized.

While this investigation was focused on using a standard IHC technique, as previously described, there are many alterations to the process of IHC that can affect staining and thus changes in fluorescence intensity over time, including the inclusion or exclusion of an antigen retrieval method and the type of detection method used. In the case of this investigation an antigen retrieval method was not utilized and the type of detection method used was a secondary antibody conjugated to a fluorophore. Observed results could differ if an antigen retrieval method or an enzyme-based detection method was used, thus results may not expand to different IHC protocols. An important future direction would thus be to carry out this investigation with various IHC protocols.

### Considerations for experimental design of fluorescence-based studies

In conclusion, primary and secondary antibody choice, primary-secondary antibody combination, and widefield versus confocal microscopy are all important factors that can contribute to the pattern of changes in fluorescence intensity over time. This can have significant effects on interpretation of images and data collected from images in contexts such as cell counting or assessment of protein expression levels. These results reinforce best practices when it comes to imaging tissue specimens. First, it is important to image shortly after staining (especially if using AF488- or AF647-conjugated secondary antibodies) in order to detect all stained cells or structures. Second, if comparing across cohorts of samples, sections from each cohort should be imaged within similar time frames after staining. This is because imaging samples at different time points post-staining does not enable accurate comparisons between experimental groups.

## Acknowledgements

Research reported in this manuscript was supported by the National Institute on Deafness and other Communication Disorders of the National Institutes of Health under Award Number R01DC018516 to C.E.J.C. and the National Institute of General Medical Sciences of the National Institutes of Health under Award Number T32GM144300. The content is solely the responsibility of the authors and does not necessarily represent the official views of the National Institutes of Health. We thank Jordan Gregory and Michael Marar for contributions to preliminary data collection and analyses.

## Notes

### Competing Interest Statement

The authors have declared no competing interest.

